# Stimulus-specific neural encoding of a persistent, internal defensive state in the hypothalamus

**DOI:** 10.1101/805317

**Authors:** Ann Kennedy, Prabhat S. Kunwar, Lingyun Li, Daniel Wagenaar, David J. Anderson

## Abstract

Persistent neural activity has been described in cortical, hippocampal, and motor networks as mediating short-term working memory of transiently encountered stimuli^1–4^. Internal emotion states such as fear also exhibit persistence following exposure to an inciting stimulus^5,6^, but such persistence is typically attributed to circulating stress hormones^7–9^; whether persistent neural activity also plays a role has not been established. SF1^+^/Nr5a1^+^ neurons in the dorsomedial and central subdivision of the ventromedial hypothalamus (VMHdm/c) are necessary for innate and learned defensive responses to predators^10–13^. Optogenetic activation of VMHdm^SF1^ neurons elicits defensive behaviors that can outlast stimulation^11,14^, suggesting it induces a persistent internal state of fear or anxiety. Here we show that VMHdm^SF1^ neurons exhibit persistent activity lasting tens of seconds, in response to naturalistic threatening stimuli. This persistent activity was correlated with, and required for, persistent thigmotaxic (anxiety-like) behavior in an open-field assay. Microendoscopic imaging of VMHdm^SF1^ neurons revealed that persistence reflects dynamic temporal changes in population activity, rather than simply synchronous, slow decay of simultaneously activated neurons. Unexpectedly, distinct but overlapping VMHdm^SF1^ subpopulations were persistently activated by different classes of threatening stimuli. Computational modeling suggested that recurrent neural networks (RNNs) incorporating slow excitation and a modest degree of neurochemical or spatial bias can account for persistent activity that maintains stimulus identity, without invoking genetically determined “labeled lines”^15^. Our results provide causal evidence that persistent neural activity, in addition to well-established neuroendocrine mechanisms, can contribute to the ability of emotion states to outlast their inciting stimuli, and suggest a mechanism that could prevent over-generalization of defensive responses without the need to evolve hardwired circuits specific for each type of threat.

## Main Text

We performed fiber photometry^16^ in VMHdm^SF1^ neurons expressing GCaMP6s^17^ in freely behaving mice during a 10s presentation of a predator (an anesthetized rat^18^) (Fig. 1a-c). We observed a rapid increase in the bulk calcium signal at the onset of rat presentation (Fig. 1d, e). Remarkably, this activity persisted for over a minute following rat removal, decaying exponentially with time constant τ_decay_ = 26.7±2.2 seconds (Fig. 1d-e, h-i; **Supplemental Video 1**). In contrast, a toy rat presented in the same manner produced a far weaker and less persistent response (Fig. 1d, e).

**Figure 1.**
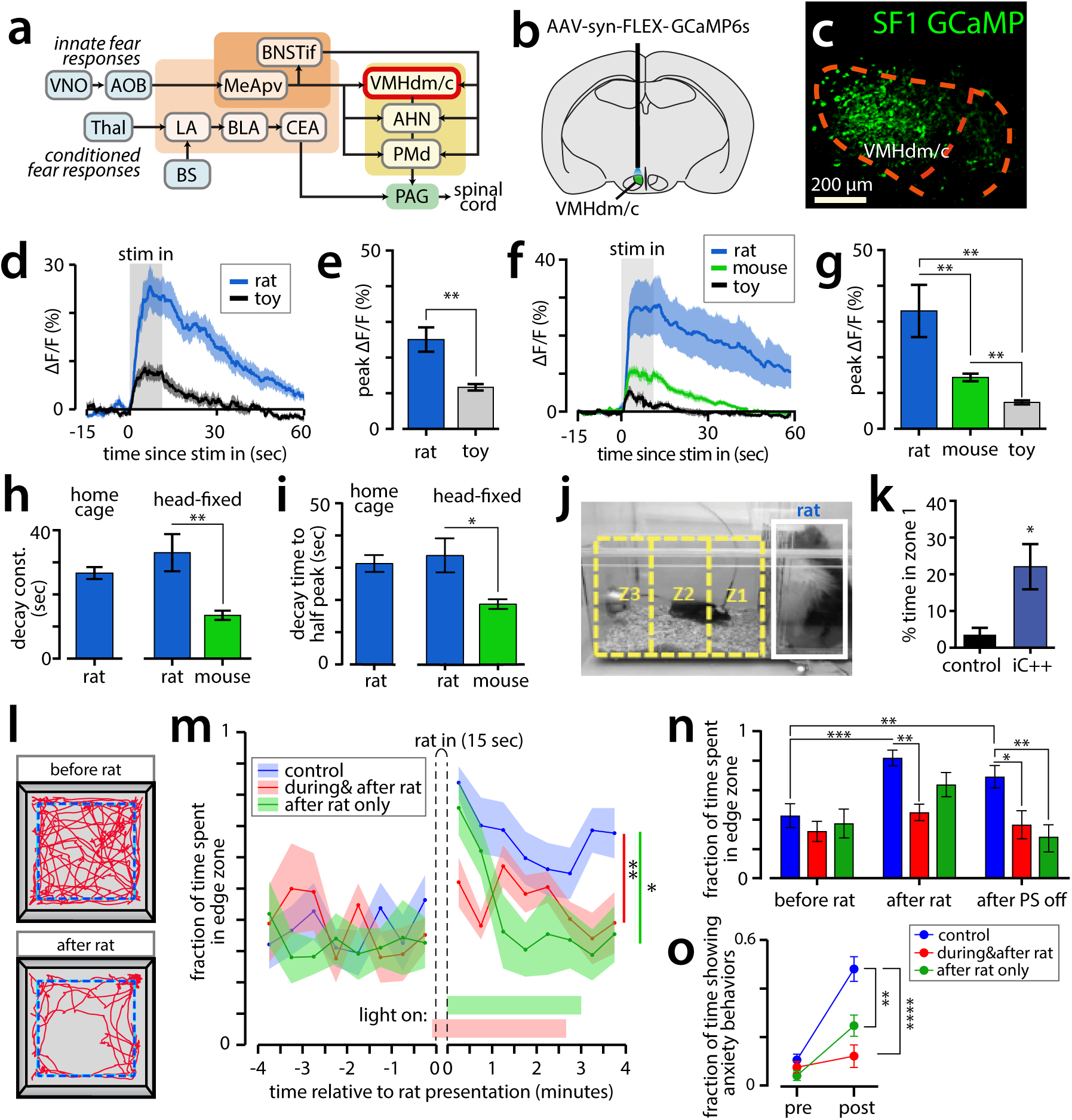
Persistent activity in VMH SF1+ neurons evoked by predatory and conspecific cues. **a**, Circuits for innate and learned fear. Abbreviations below. **b**, Site of fiber photometry in VMHdm/c (green). **c**, GCaMP6s expression in VMHdmSF1 neurons. **d**, Activity of SF1+ neurons in freely moving mice exposed to a live or toy rat for 10 seconds (gray shading). (n = 4 mice; mean ± SEM). **e**, Peak activity from (d) (n = 4 mice; mean ± SEM). **f**, Responses of VMHdmSF1 neurons in head-fixed mice to rat, mouse, or toy rat (n=8 mice; mean ± SEM). **g**, Peak activity from (f) (n = 8 mice; mean ± SEM). **h, i**, Decay constants (h) or time to half-peak (i) of activity in freely moving (“home cage”) or head fixed mice (mean ± SEM; home cage n = 4; head-fixed n = 8). **j**, Home cage rat exposure assay. **k**, Percent time in zone 1 during 3-minute rat presentation (PS; n=7 control mice; n=7 iC++ mice; mean ± SEM). **l**, Tracking of mouse in open field rat exposure assay, blue line marks “edge zone”. **m**, Fraction of time spent in edge zone. Colored horizontal bars denote PS periods (n = 12 control mice, n = 6 iC++ mice with PS during+after rat; n = 6 iC++ mice with PS after rat only. * p < 0.05; ** p< 0.01, repeated measures ANOVA test. Data are mean ± SEM.) **n**, Mean time in edge zone, times defined in Methods. **o**, Expression of anxiety behaviors (see Methods) before vs after rat exposure. (mean ± SEM). VNO - vomeronasal organ, AOB - accessory olfactory bulb, MeApv –posterioventral medial amygdala, BNSTif – interfas-cicular part of bed nucleus of the stria terminalis, VMHdm – dorosmedial ventromedial hypothalamus, AHN – anterior hypothalamic nucleus, PMd - dorsal premammillary nucleus, PAG - periaqueductal gray, Thal – thalamus, LA – lateral amygdala, BS – brain stem, BLA – basolateral amygdala, CEA – central amygdala.

To better control the timing and location of stimulus presentation, we repeated this experiment using a head-fixed preparation. The magnitude, decay constants and specificity of VMHdm^SF1^ persistent responses were comparable to those measured in freely behaving mice (Fig. 1f-i; Extended Data. 1). As discussed below, this persistence is unlikely to be due simply to slow GCaMP decay kinetics (Fig. 2 and Extended Data Fig. 5). VMHdm^SF1^ neurons also responded to rat urine alone (Extended Data Fig. 2), consistent with earlier studies^19,20^.

**Figure 2.**
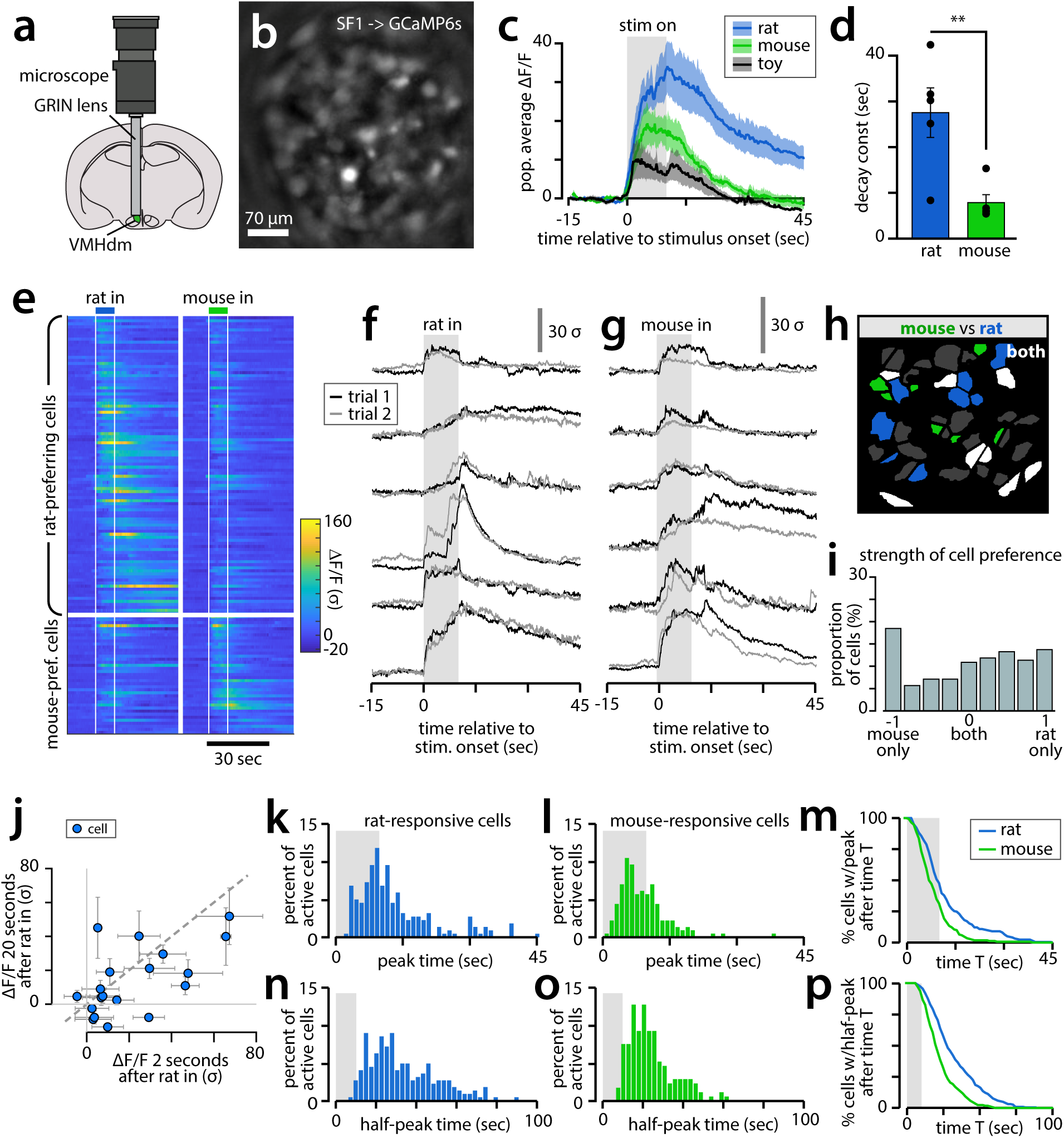
Microendoscopic imaging reveals persistent activity at the single cell level. **a**, Microendoscopic imaging in VMHdm/c. **b**, Field of view in an imaged mouse. **c**, Mean population response of imaged neurons to each stimulus (n = 2 trials/mouse from 5 mice, mean ± SEM). **d**, Fit decay constants of population response to rat and mouse (n=5 mice, mean ± SEM). **e**, Rat- and mouse-responsive neuron responses (from n=5 mice, mean over 2 trials). **f**, Example cells responding to rat in one imaged mouse on two repeated trials. **g**, Example cells responding to mouse on two repeated trials (same mouse as (f)). **h**, Example spatial map of cells responsive to rat, mouse, or both (white). **i**, Histogram of cell tuning preference for rat vs. mouse. Cells at ± 1 respond exclusively to rat or mouse, respectively; cells at 0 (“both”) respond equally to both stimuli (n = 219 cells from 5 mice across 3 days of imaging). **j**, Scatterplot comparing cell responses at 2 vs 20 seconds after rat introduction, in one example mouse. **k**, Peak time for rat-responsive cells (n = 202 rat-responsive cells from 5 mice across 3 days of imaging). **l**, Peak time for mouse-responsive cells (n = 160 mouse-responsive cells from 5 mice across 3 days of imaging). **m**, Fraction of cells with peak after time T. **n**, Half-peak time for rat-responsive cells (n same as k). **o**, Half-peak time for mouse-responsive cells (n same as l). **p**, Fraction of cells with half-peak later than time, legend as in (m).

To investigate whether VMHdm^SF1^ neurons were also active during persistent defensive behaviors, we devised a novel rat exposure assay in an open arena. Following a ten-minute period of habituation to the arena by the mouse, an awake rat in a cage was presented to the mouse for 15 seconds, and then removed. After rat exposure, mice exhibited thigmotaxis, an indication of increased anxiety^21^, lasting for minutes (Fig. 1l, m; **control; blue line**). Thigmotaxis was not observed if the mouse was introduced to the arena following, rather than before, rat presentation, indicating that it is unlikely to be due to residual rat-derived odors (Extended Data Fig. 3a-c). Persistent thigmotaxis could also be evoked by brief optogenetic stimulation of VMHdm^SF1^ neurons (ref ^11^ and Extended Data Fig. 3d, e)). Fiber photometry confirmed that VMHdm^SF1^ neurons were strongly and persistently activated for minutes following rat presentation in the arena (Extended Data Fig. 4c-e), with kinetics well correlated with behavior (Extended Data Fig. 4f-i).

We next tested a requirement for VMHdm^SF1^ neurons in the rat-evoked increase in thigmotaxic behavior, using the light-gated chloride channel iC++^22^ to reversibly silence these cells. First, we confirmed that iC++ inhibition of VMHdm^SF1^ neurons in a home cage rat exposure assay increased the time the mice spent in close proximity to the rat (Fig. 1j, k), phenocopying genetic ablation of these neurons^11^. Next, we repeated the open field rat exposure test while photo-inhibiting VMHdm^SF1^ neurons continuously for a 3 min period, either starting five seconds prior to rat introduction, or immediately following rat removal (Fig. 1m; **“light on,” red vs. green bars, respectively**).

When silencing was initiated five seconds prior to rat introduction, no significant rat-induced increase in thigmotaxis was observed (Fig. 1m-o; red plots). Importantly, when iC++ photostimulation was initiated only after rat removal, mice showed an initial increase in thigmotaxis, but quickly returned to their pre-rat baseline behavior (Fig. 1m-o, green plots; Extended Data Fig. 4a). These data indicate that VMHdm^SF1^ neuronal activity is essential for maintaining a persistent defensive behavioral response to a predator.

To investigate the neural dynamics underlying the rat-evoked persistent state, we next performed microendoscopic imaging^23,24^ of VMHdm^SF1^ neurons (Fig. 2a-b). Head-fixed mice were presented for ten seconds each with a pseudorandomized set of stimuli including a rat, toy rat, and conspecific male, on each of three days of imaging (n=5 mice, 187.3±8.1 cells imaged per day, 78 cells tracked across days.) While VMHdm^SF1^ neurons, as a population, showed persistent activation following rat presentation (Fig. 2c-d), individual VMHdm^SF1^ neurons showed diverse but reproducible stimulus-evoked dynamics (Fig. 2e-g). Although many cells showed activation from stimulus onset followed by slow decay, other cells reached their peak activation only after stimulus removal. Thus the slow, monotonic decay of the population response reflects a diverse, time-evolving pattern of activity at the level of individual cells (Fig. 2j-p). Application of a spike-deconvolution algorithm^25^ indicated that the persistent ∆F/F signal likely reflects persistent underlying spiking activity (Extended Data Fig. 5). Strikingly, the rat and conspecific activated distinct subpopulations of VMHdm^SF1^ neurons with only moderate overlap (Fig. 2h, i).

In rodents, nonvolatile odor cues can activate neurons of the vomeronasal organ (VNO) for several seconds following inhalation^26^. It was therefore possible that the persistent activity we observed in VMHdm^SF1^ neurons reflected persistence of the stimulus signal. If this were the case, we would expect VMHdm^SF1^ neurons to show only transient responses to time-resolved stimuli, such as purely visual or auditory stimuli. VMHdm neurons are not activated by (Extended Data Fig. 6), or required for^11^ defensive responses to, an overhead visual threat stimulus^27^, and have not previously been shown to respond to auditory stimuli. We therefore imaged VMHdm^SF1^ neuron activity in response to an auditory stimulus that evokes defensive behaviors in mice^28,29^: a series of ultrasonic sweeps (USS) in the frequency range typical of rat distress vocalizations^30^ (Methods).

The USS strongly activated VMHdm^SF1^ neurons, and this activation persisted on a similar time scale as that following rat exposure, after stimulus termination (Fig. 3a-b). Underlying this population response, individual USS-responsive VMHdm^SF1^ neurons showed diverse dynamics, as was observed for the rat and mouse stimuli (Fig. 3c-f). The USS activated VMHdm^SF1^ neurons were spatially intermingled with the rat (and mouse)-responsive populations, and overlapped them at a frequency equal to that expected by chance (Fig. 3g-j; Extended Data Fig. 7). This suggests that these populations are randomly distributed, but reproducibly activated by a given stimulus. Of the neurons showing a significant response to at least one of our five test stimuli (74.4%), 43.6% responded to only one stimulus, and over 70% of cells responded to ≤ 2 of the five tested stimuli (Fig. 3k). Most stimulus-responsive VMHdm^SF1^ neurons were excited, although a small number showed stimulus-evoked inhibition (Fig. 3c-f, l). Principal component analysis (PCA) indicated that in addition to stimulus identity, VMHdm^SF1^ neurons may encode an additional feature common to multiple stimuli (Fig. 3o, y-axis). Since this feature has higher values on initial vs. later trials, it may relate to novelty or salience. A 5-way Naïve Bayes decoder was able to predict stimulus identity in held-out trials across three days of imaging with above-chance accuracy (Fig. 3m). Importantly, despite the gradual decay in population activity following stimulus offset, stimulus identity could still be decoded with above chance accuracy for tens of seconds (Fig. 3n, Extended Data Fig. 8). Thus, the VMHdm^SF1^ population response can encode the identity of the presented stimulus, even after its removal.

**Figure 3.**
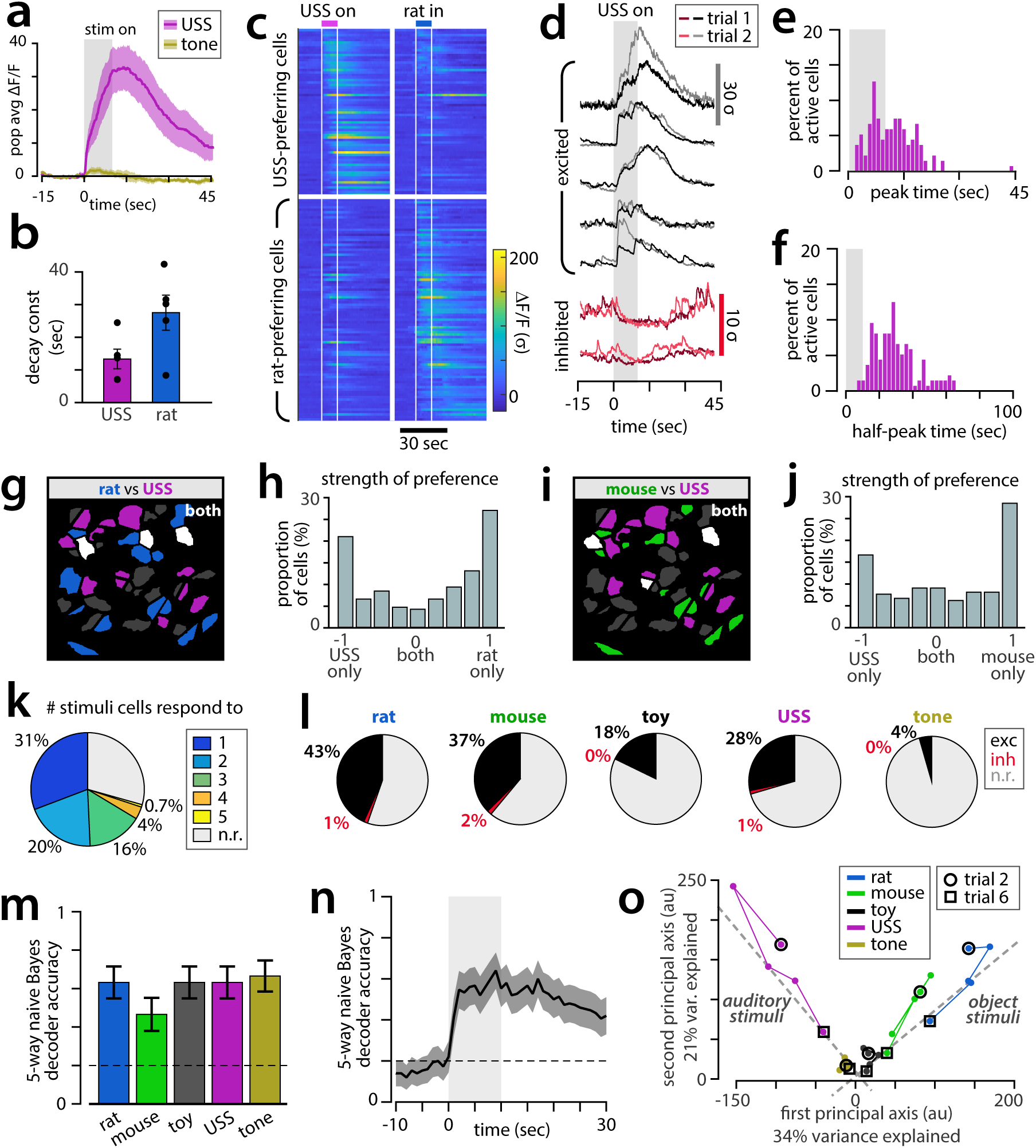
SF1+ neurons respond to a threatening auditory stimulus, and encode stimulus identity. **a**, Mean VMHdmSF1 population response to USS and tone (n = 5 mice, mean ± SEM). **b**, Fit decay constants of population response to USS (rat reproduced from Fig2d for comparison). **c**, Example cells responding to USS in one imaged mouse, on two repeated stimulus presentations (dark trace: 1st presentation; light trace: 2nd presentation). Black = excited cells, red = inhibited cells. **d**, Response peak time of USS-responsive cells (n = 133 USS-responsive cells from 5 mice across 3 days of imaging). **e**, Response half-peak time for USS-responsive cells (n same as (d)). **f**, Example spatial map of cells responsive to rat, USS, or both (white). **g**, Histogram of cell tuning preference, rat vs USS (n = 216 cells from 5 mice across 3 days of imaging). **h**, Example spatial map of cells responsive to mouse and USS. **i**, Histogram of cell tuning preference, mouse vs USS (n = 219 cells from 5 mice across 3 days of imaging). **j**, Percent of cells responding to zero to five out of five stimuli. **k**, Percent of cells excited or inhibited by each stimulus. **l**, Accuracy of a 5-way Naïve Bayesian decoder for stimulus identity, trained on six trials across three days of imaging (n=5 mice; mean ± SEM). **m**, Decoder accuracy as a function of time. **n**, Principal component analysis (PCA) of time-averaged population responses, pooled across mice, from 5 trials per stimulus across 3 days of imaging.

We next used computational modeling to investigate the space of mechanisms that can account for persistent activity in VMHdm^SF1^ neurons, incorporating features that have been proposed in other systems^2,3,31-36^. We evaluated four classes of models (Fig 4a1-4) in terms of their ability to capture two main features of VMHdm^SF1^ neural activity: it changes over time, and it is stimulus-specific. To achieve persistent activity with time-evolving dynamics that aligned with our experimental observations, it was necessary to combine two elements: inhibition-stabilized recurrent connectivity and slow excitation on a time scale of several seconds (mediated by, e.g., peptidergic transmission^37^); we call the resulting class of models peptidergic recurrent neural networks (pRNNs) (Fig. 4a3). The importance of feedback inhibition in our model suggests the existence of an inhibitory population that is activated by threatening stimuli; a likely candidate are GABAergic neurons in the surrounding DMH^38,39^.

**Figure 4.**
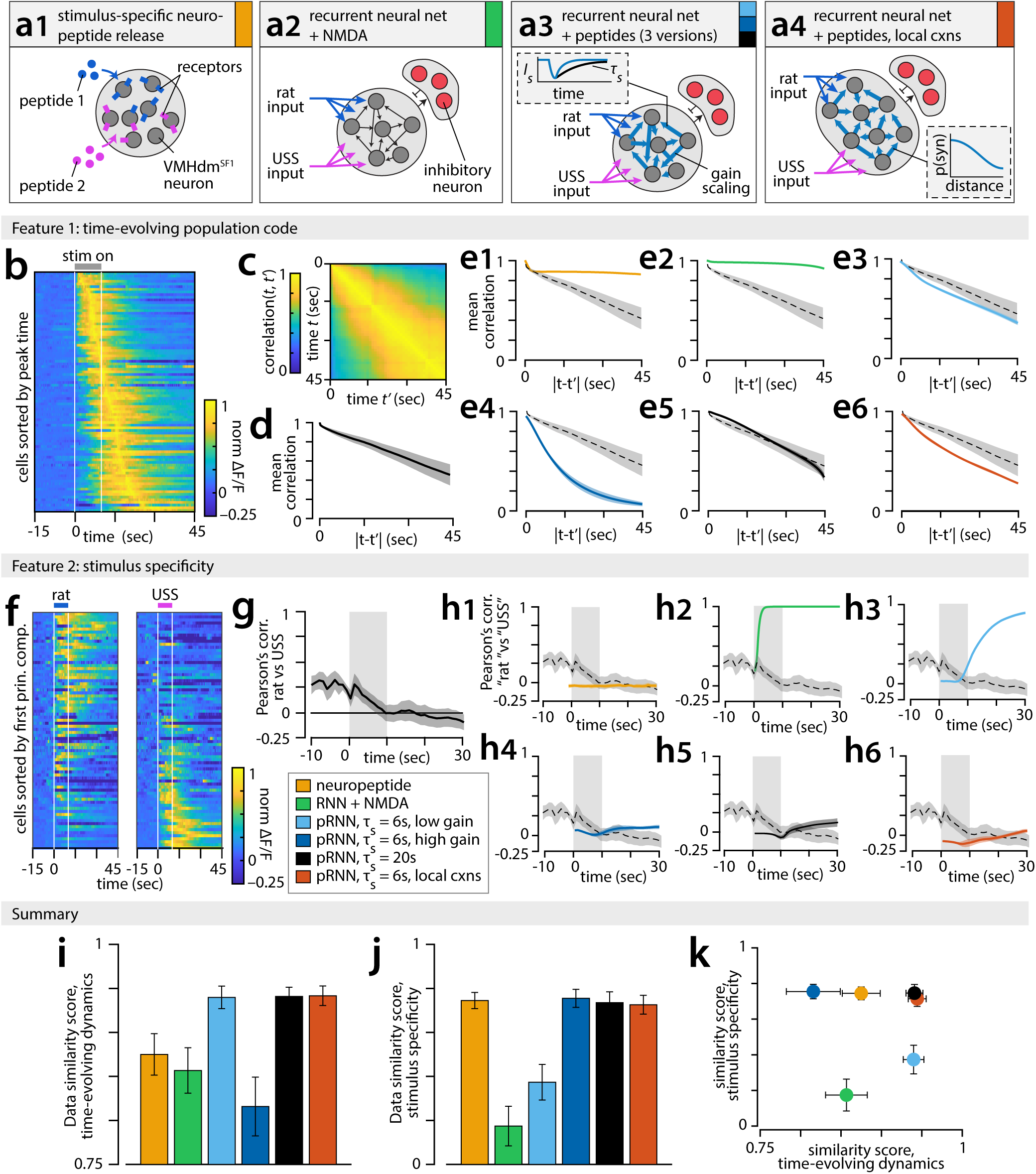
Data constrains set of possible mechanisms for persistent neural activity. **a** Six tested models of persistent activity. **a1** slow-acting neuropeptide activation, no connectivity between model neurons. **a3** recurrent excitation in a randomly connected network with fast inhibitory feedback; persistence maintained via NMDA channels, time constant of NMDA excitation is τ_s_ = 200ms. **a3** as in **a2**, replacing NMDA with slow peptidergic excitation (light/dark blue, τ_s_ = 6sec; black, τ_s_ = 20sec), tested for different strengths of recurrent synapses (“gain scaling”, light/dark blue). **a4** as in **a3**, but with “local connectivity” in which probability of a synapse *p(syn)* between neurons decreased with distance (inset). **b** Trial-averaged, normalized ΔF/F traces from USS-responsive neurons, sorted by time of response peak. **c** Autocorrelation matrix of USS-evoked population activity. **d** Time-averaged autocorrelation of USS-evoked population activity (n=5 mice, mean ±SEM). **e** Autocorrelation as in **d**, for each model; model identity indicated by line color (legend in panel **h**), dashed line = data (n=10 repeat simulations, mean ± SEM). **f** Trial-averaged ΔF/F traces from rat-or USS-responsive neurons, sorted by projection on first principal component. **g** Pearson’s correlation between rat-vs USS-evoked population activity as a function of time. **h** Pearson’s correlation between simulated “rat” and “USS” inputs to each model, dashed line = data (n=10 repeat simulations, mean ± SEM). **i** “Similarity score” (see Methods) of models vs. data, summarizing plots in **(e)**. (n=5 mice, mean ± SEM). **j** Similarity score of model vs data, summarizing plots in **h**. (n=5 mice, mean ± SEM). **k** scatter plot of **i-j**.

To compare the time-evolving population code of VMHdm^SF1^ neurons (Fig4b) with the dynamics of these models, we computed the autocorrelation matrix of stimulus-evoked activity for all values of *t* and *t’* between 0 and 45 s after stimulus onset(Fig 4c), and summarized it by computing the mean correlation over all neurons as a function of ∆t (*t* – *t’*) (Fig 4d). Several versions of our pRNN models showed autocorrelation dynamics similar to observed dynamics (Fig 4e). Importantly, models that used slow peptidergic transmission alone (Fig 4a1), or RNNs with NMDA-mediated transmission^40^ (Fig 4a2) could not match the diverse responses we observed in VMHdm^SF1^ neurons (Fig. 4e1-2).

The stimulus specificity of VMHdm^SF1^ responses (Fig 4f) was quantified as the time-evolving Pearson’s correlation between rat-vs. USS-evoked activity (Fig 4g). Maintaining stimulus specific representations during persistence could only be achieved in the pRNN by increasing the gain (strength) of excitatory synapses^41,42^ (Fig. 4a3, h3-4), but this came at the cost of failing to match the autocorrelation dynamics of observed activity (**Fig e3-4**). However, both main features of the data could be matched if the synaptic time constant of peptidergic excitation was further increased from 6 to 20 seconds (Fig 4a3,h5,e5). Alternatively, both features could be matched by assuming that the probability of synapse formation between a pair of model neurons decreased slightly with the distance between them, and imposing a mild differential spatial bias on the responses to different stimuli (rat vs. USS) (Fig 4a4,h6,e6, ED Fig 9). Imposing further network structure produced too much segregation between stimulus representations, compared to the overlap observed (**ED Fig 9**). We summarized the performance of our models by creating a pair of “data similarity scores” quantifying model similarity to data in terms of time-evolving dynamics and stimulus specificity (Fig 4i-k).

Persistent defensive states are typically attributed to neuroendocrine mechanisms, such as activation of the HPA axis^7–9^. Here we provide the first evidence that persistent neural activity can contribute causally to such persistent internal states. VMHdm^SF1^ activity may also activate longer-lasting neuroendocrine processes, as stimulation of VMHdm^SF1^ neurons elevates serum cortisol levels^11^. However unlike circulating hormones, persistent activity in VMHdm^SF1^ neurons is stimulus-specific, and may thereby prevent over-generalization of defensive responses. The observed persistent activity can be modeled best by recurrent excitatory networks incorporating fast feedback inhibition and slow peptidergic transmission^43^. VMHdm^SF1^ neurons are exclusively glutamatergic, densely interconnected^44^, and express several neuropeptides as well as neuropeptide receptors^45^, properties consistent with optimal model features. While our data do not exclude a role for interconnected structures^10,46^ in establishing persistent activity in VMHdm, they demonstrate that hypothalamic neuronal population dynamics contribute to the persistence of emotional behaviors.

## ACKNOWLEDGMENTS

We thank H. Inagaki, M. Meister, L.F. Abbott, U. Rutishauser, and members of the Anderson lab for helpful comments on the manuscript, T. Davidson and K. Deisseroth for help implementing fiber photometry, R. Remedios for help with miniscope imaging, X. Da, J.S. Chang and C. Kim for technical assistance, X. Da and C. Chiu for laboratory management and G. Mancuso for administrative support. This work was supported by NIH Grant R01 MH112593. K99 MH117264 to A.K. and a Helen Hay Whitney Foundation Postdoctoral Fellowship to L. L. D.J.A. is an Investigator of the Howard Hughes Medical Institute.

## METHODS

### Animals

All experimental procedures involving the use of live animals or their tissues were performed in accordance with the NIH guidelines and approved by the Institutional Animal Care and Use Committee (IACUC) and the Institutional Biosafety Committee at the California Institute of Technology (Caltech). SF1-Cre mice were obtained from Dr. Brad Lowell ^1^ and maintained as heterozygotes in the Caltech animal facility as described previously; the SF1-Cre line is also available from the Jackson Laboratory (Stock No: 012462). An account of the specificity of SF1-Cre expression within VMH and characterization of neurons labeled by Cre-expression can be found in ^2^. Male mice, heterozygotes or their wild-type littermates aged between 8 to 20 weeks were used in this study. All mice were housed in ventilated micro-isolator cages in a temperature- and humidity-controlled environment under a reversed 12 hour dark-light cycle, and had free access to food and water. Mouse cages were changed weekly on a fixed day on which experiments were not performed. Long-Evans rats (for use as predators) were obtained from Charles River at 2-3 months of age, and raised to 5-10 months in the Caltech animal facilities.

### Virus

AAV1.Syn.Flex.GCaMP6s.WPRE.SV40 (CS1113) was obtained from the Penn Vector Core. AAV5.EF1a.DIO.iC++.eYFP and AAV2.EF1a.DIO.hChR2.eYFP.WPRE.pA were obtained from the University of North Carolina Vectors Core.

### Surgery

Mice 8-20 weeks old were anesthetized with 5% isoflurane and mounted in a stereotaxic apparatus (Kopf Instruments). 1% - 1.5% isoflurane was used to maintain the anesthesia throughout the surgery procedure. An incision was made to exposure the skull and small craniotomies were made dorsal to each injection site with a stereotaxic mounted drill. Virus suspension (~600 nl) was injected to the VMHdm/c (ML +/- 0.5, AP −4.65, DV −5.6) at a rate of 60 nl/minute using a pulled glass capillary (~40 µm inner diameter at tip) mounted in a nanoliter injector (Nanoliter 2000, World Precision Instruments) controlled by a four channel micro controller (Micro4, World Precision Instruments). Capillaries were kept in place for 10 minutes following injections to allow the adequate diffusion of virus solution and to reduce the virus backflow during capillary withdraw.

For fiber photometry, a custom-made unilateral fiber cannula (400 µm in core diameter, 0.48 NA, Doric Lenses) was implanted after virus injection (ML +/-0.4, AP −4.65, DV −5.4). Metabond (Parkell) and dental cement (Bosworth) were applied to secure the implanted ferrule and cover the exposed skull. For optogenetics, a custom-made bilateral fiber cannula aimed 500 µm above each injection site (200 µm in core diameter, 0.37 NA, Doric Lenses) was implanted and held in place with Metabond and dental cement.

Surgery for microendoscopic imaging was performed as previously described^3^. Briefly, we first performed a series of titration experiments of the original viral stock, to determine the virus concentration at which the brightest cytoplasmic but non-nuclear GCaMP6s expression could be observed in slices of fixed brain tissue of the injected mice 4 weeks after injection. The optimal viral dilution was then used to inject mice for *in vivo* imaging as described above. 2-3 weeks after viral injections, mice were implanted with a graded-index (GRIN) lens (diameter − 0.5 mm, length − 8.4 mm, catalogue #1050-002212, Inscopix) using a supporting device (Proview Implant Kit, cat# 1050-002334, Inscopix). The implantation depth of the lens was determined based on the live visualization of (anesthetized) neural activity as the lens was inserted. Metabond was used to stabilize the lens, and Kwik-Sil sealant (World Precision Instruments) was used to cover the lens surface. After another 2-3 weeks, mice were anesthetized for placement of a microendoscope baseplate (cat# 1050-002192, Inscopix) and a baseplate cover (catalogue #1050-002193, Inscopix) was used to protect the lens when not in use. Five out of twenty implanted animals were selected for *in vivo* imaging studies based on clarity of cytoplasmic GCaMP6s expression.

### Stimuli Presentation

Stimuli were presented either in the mouse’s home cage or in a head-fixation set up. In the home cage, a hand-held anesthetized rat weighing 400-600 g was brought in close proximity to the mouse. A stuffed toy rat of approximately the same size as the live rat was used as a control. For the head-fixed preparation, the mouse was placed on a plastic running wheel (15.5 cm diameter) and stabilized by the head-plate (World Precision Instruments, Catalogue #503617) with a custom made tethering system. Animals were habituated to the head-fixation setup for 1 hour each day for 2-3 days before experiments began. Physical stimuli (an awake behaving rat, a conspecific BALB/6 male mouse or a toy rat) were each presented inside a small wire mesh cage, which was held by the experimenter in front of the experimental mouse. Auditory stimuli were presented at 85 dB SPL from above the animal. The ultrasound stimulus (USS) consists of repeated 100 ms frequency sweeps from 17-20 kHz, as described previously ^4^. A pure tone of 2 kHz was used as a control. Rat urine was collected in-house and kept at 4°C for up to two weeks. A cotton swab soaked with 100 µl of rat urine or water was presented in front of the experimental mouse. 500 ms looming stimulus was displayed on an overhead screen above the mouse home cage 10 times with 500 ms inter stimulus interval. All stimuli were pseudo-randomized and presented for 10 seconds unless otherwise clarified, with an inter-trial interval of at least five minutes. For microendoscopic imaging, two trials for each stimulus were presented on each of three consecutive days.

### Optogenetic manipulation

Optogenetic experiments were performed as described in ^2^. Animals were briefly anaesthetized by isoflurane to connect the fiberoptic patch cord to the bilateral implanted optic cannula (Doric Lenses). Mice were then allowed to recover for at least 15-20 minutes in their home cage before being transferred to the behavioral testing room. Light for both iC++ and ChR2 activation was delivered via a 473nm laser (Shanghai Laser) controlled by a signal generator (A-M systems, isolated pulse stimulator). Laser intensity was calibrated at the distance of 0.5 mm below the implanted fiber tip. 3 minutes continuous light was used for iC++ activation; 10 seconds (20 Hz, 20 ms pulse width) pulse trains was used for ChR2 activation.

### Home cage rat exposure assay

The mouse home cage was placed into a custom made testing apparatus (35 x 40 x 40 cm), and video of behavior was collected from a side-view camera. After a 6 minute baseline, a predator rat in a cage with a mesh wall (10 x 20 x 35 cm) was placed at one end of the mouse home cage. Ethovision XT software was used to track mouse position and quantify time spent in proximity to the rat.

### Open field rat exposure assay

The mouse was placed in a plastic open top arena (50 x 50 cm, 30 cm walls), with behavior captured using an overhead mounted camera. Following a 10 minute baseline, a rat held in a cage with a mesh wall was held in close proximity to the mouse for 15 seconds, and then removed. Behavior of the mouse was then recorded for an additional 6 minutes. For behavior quantification, Ethovision tracking data was segmented into 30-second chunks, and percent of time spent in the “edge zone” (within 4cm of arena walls) was quantified. For bar graphs in Fig 1n, we define before rat = average over a window from −1 to 0 min relative to rat presentation, after rat = average from 0-1 min after the rat was removed, and after PS off = average from 3-4 min after rat was removed. Anxiety behaviors for Fig 1o were defined as thigmotaxis, immobility, and jumping (escape attempts) and were manually annotated at 30Hz. Pre and post windows correspond to −3 to 0 and 0 to 3 min, respectively, relative to rat presentation.

### Fiber photometry data acquisition and processing

Fiber photometry was performed as described in ^5^. Briefly, two LEDs modulated at different frequencies (490 nm and 405 nm, Thorlabs) were used to excite GCaMP6s-expressing neurons via implanted optical fiber. Excitation light at 490 nm activates GCaMP6s in a calcium-dependent manner, while excitation at 405 nm activates GCaMP6s in a calcium-independent manner, thus the 405nm signal can be used to control for bleaching and movement artifacts in the 490nm channel. A photometer (Newport Femtowatt Photoreceiver) received GCaMP6s fluorescent signals, and custom-designed software separated the signals generated by the two LEDs. The output power of both LED was set between 30-50 µW at the fiber tip to obtain an optimal baseline fluorescence without photobleaching.

To calculate ΔF/F of the 490nm signal, we normalized it to the 405nm baseline as in ^5^. The 405nm signal was scaled to match the amplitude of the 490nm signal using linear regression, and ΔF/F computed as (490nm signal – scaled 405nm signal) / (scaled 405 nm signal).

### Microendoscopic imaging data acquisition and processing

We used a head-mounted miniaturized microscope (nVista, Inscopix) for calcium imaging. Pilot experiments were done to identify imaging parameters that produced the clearest signal to noise ratio while limiting photobleaching. All mice except one were recorded at 11 Hz with 90.0ms exposure time, 10-20% LED illumination and 1.5 − 2.5× gain; the remaining mouse was imaged at 20Hz with 50ms exposure time. A custom-built system was used to synchronize the cameras for behavioral recordings and devices for neural recordings and stimuli delivery.

Imaging frames were spatially downsampled by a factor of two in the X and Y dimensions, and spatially high-pass filtered with a cutoff spatial frequency of 40 µm. All frames collected over the course of a single day were then concatenated into a single stack and registered to each other to correct for motion artifacts using a rigid-body transformation (TurboReg plugin for ImageJ). Single cell Ca^2+^ activity traces and spatial filters were extracted from the registered movie using CNMF-E^6^. Extracted traces and ROIs were manually screened to remove neuropil or other non-neuronal signals. The cleaned set of cells were then registered across three consecutive days of imaging as described in^3^. Briefly, all extracted spatial filters from a given day of imaging were added to create a cell map, and intensity-based image registration was used to identify a pair of rigid-body transformations to align the day 1 and 3 maps to the day 2 maps. Overlapping triplets of spatial filters for the three days were identified by finding cells on day 1 and 3 with the smallest Euclidean distance to each day 2 cell. All identified triplets were then manually screened for accuracy. Roughly half of all cells could be registered across all three days of imaging.

### Spike inference

Spiking of Sf1^+^ neurons was estimated from extracted GCaMP fluorescence traces using the constrained deconvolution approach of ^7^. Calcium transients were modeled as a first-order autoregressive (AR) process, and with the exception of the AR coefficient γ (GCaMP decay rate), model parameters were fit as in ^7^ to all data from each cell on a given day of imaging. The value of γ was selected to give a GCaMP half-decay time of two seconds, following the expected decay kinetics of GCaMP6s in high firing rate conditions^8^. This is a conservative estimate (other reported half decay times of AAV-GCaMP6s range from 510ms to 1.8sec^9^) to obtain an approximate upper limit on the contribution of slow GCaMP dynamics to the observed persistence of SF1^+^ population activity.

### Units of neuronal activation

Stimulus-evoked responses of SF1^+^ neurons are reported in units of baseline standard deviation, σ, defined as the standard deviation of observed fluorescence in a 30-second pre-stimulus baseline.

### Fitting decay constants in fiber photometry and population average microendoscope data

The stimulus-evoked response of the SF1^+^ population could be well fit by a difference of exponentials of the form *K*(*t*) = *A* ∙ (*τ*_*decay*_ − *τ*_*rise*_)^−1^(*e*^−*t*/*τ*rise^ − *e*^−*t*/*τ*decay^), where *t* is time in seconds, and *A*, *τ*_*decay*_, and *τ*_*rise*_ are fit parameters characterizing the amplitude and kinetics of the response. Values of, *τ*_decay_, and *τ*_rise_ were fit for each trial to minimize the mean squared error between *K*(*t*) and the SF1^+^ population response over a 30 second window following the start of stimulus presentation, using the *fminunc* function in Matlab.

### Identifying stimulus-responsive cells

Some analyses, such as calculation of time to peak or half-peak strength of stimulus preference, were performed only on cells that showed a significant response to the stimulus or stimuli in question. For these analyses, we defined a pre-stimulus baseline as the ΔF/F in a ten-second window prior to stimulus presentation, and defined responsive cells as those neurons for which the average ΔF/F value for any one-second window in the first 30 seconds after stimulus presentation was more than four standard deviations above the mean of baseline activity on that trial. Only cells that passed this criterion on both trials within a day were included for analysis.

### Finding peak and decay to half-peak times

The time of the peak population response was defined as the first time (relative to start of stimulus presentation) the population ΔF/F passed 95% of its maximum observed value on a given trial. The time to decay to half-peak was defined as the last time the population ΔF/F was above 50% of its maximum value relative to pre-stimulus baseline. This analysis was performed separately for each cell on each day of imaging, using only stimulus-responsive cells (see above); values were averaged across two repeated stimulus presentations.

### Strength of cell preference

The strength of cell preference for either of a pair of stimuli (Fig 2g and Fig 3i,k) was defined as |s_*a*_ − s_*b*_|/(|s_*a*_| + |s_*b*_|), where s_*a*_ and s_*b*_ are the average ΔF/F of that cell in a 45-second window following stimulus onset, for stimulus pair *a* vs *b* (eg rat vs USS). This analysis was performed separately for each cell on each day of imaging, using only cells that responded to either one (or both) of the two stimuli, as defined above; values of s_*a*_ and s_*b*_were averaged across two repeated stimulus presentations.

### Decoder analysis

Stimulus identity was decoded from the activity of all SF1^+^ neurons that could be reliably tracked across three days of imaging, using a two-class or five-class cross-validated Naïve Bayes decoder (*fitcnb* in Matlab). Bar plots of decoder accuracy (Fig 3n) and confusion matrix (**ED 7a**) were generated using held-out test data for a five-class Naïve Bayes decoder trained on the time-averaged responses of imaged neurons in a window from 30 seconds before to approximately 60 seconds after stimulus presentation. Time-evolving plots of decoder accuracy (Fig 3o, ED 8b-c) were constructed by training a separate cross-validated decoder on the time-averaged activity of imaged neurons in a one-second window, for each one-second window from 10 seconds before to 30 seconds after stimulus presentation. Decoder performance is reported as the average prediction accuracy on held-out test data; chance accuracy is 1/2 for the two-class decoder and 1/5 for the five-class decoder.

### Stimulus-evoked autocorrelation

We constructed the standard correlation matrix C of VMHdm^SF1^ cell activity, defined as the pairwise correlation coefficient between all columns of a neurons x time matrix of imaged activity. Values in C were averaged across each trial for a given stimulus over three days of imaging, and then averaged across n=5 imaged mice, for all imaging frames from zero to 45 seconds relative to the onset of stimulus presentation (imaging framerate was 11Hz). The mean correlation for lag Δt was computed by averaging C(x,x+Δt) for all x between 0 and 45-Δt seconds.

The same calculation was used for simulated data, with correlations computed every 10 simulation timesteps (10ms). To make values comparable to the experimental data, model cell spikes were convolved with a pair of exponential filters with time constants of 0.5 seconds and 1.5 seconds, simulating the kinetics of the GCaMP6s response.

### Rat/USS Pearson’s correlation

The Pearson’s correlation between rat and USS responses was computed for each mouse using the trial-averaged response of all neurons on the first day of imaging. Pearson’s correlation was computed between the vectors of population activity from 10 seconds before to 30 seconds after stimulus onset, sampled at 11Hz (acquisition frequency).

For simulated data, the “rat” and “USS” inputs were assumed to be excitatory inputs to a randomly selected fraction of neurons in the model (temporal structure of stimulus and percent of neurons receiving input specified below for each model). Pearson’s correlation between these two stimuli was computed across all model cells that fired 10 or more spikes across the two stimuli. GCaMP6s kinetics were simulated as for the stimulus-evoked autocorrelation analysis.

### Neuropeptide model

For this model we assumed that VMHdm^SF1^ neuron dynamics were determined entirely by long-lasting peptidergic input, and that there were no recurrent connections between neurons within VMHdm. Given a model population of N = 1000 neurons, we assumed that a random 10% of neurons received peptidergic input for any given stimulus. For cells receiving stimulus-evoked input, we modeled the firing rate *r*_*i*_(*t*) of neuron *i* as *r*_*i*_(*t*) = *g* ∙ *p*_*i*_(*t*), where *g*~*U*(0,60) sets the strength of excitatory peptidergic input onto neuron *i* and *p*_*i*_(*t*) is a stimulus-evoked peptide-mediated excitatory current. Dynamics of *p*_*i*_(*t*) evolve as 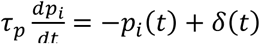, where *δ*(*t*) is the delta function and *τ*_*p*_ = 25 sec, the decay time constant that sets the duration of peptidergic excitation, was set to match the observed decay time constant of the population average VMHdm^SF1^ response. To simulate spiking, the firing rate *r*_*i*_(*t*) was used to set the instantaneous rate constant of a non-homogeneous Poisson process with a simulation timestep of dt = 1ms.

### Spiking recurrent neural network model + NMDA

We constructed a model population of N = 1000 standard current-based leaky integrate-and-fire neurons, in which each neuron has membrane potential *x*_*i*_ characterized by dynamics 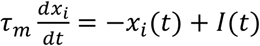, where *τ*_*m*_ = 20ms is a membrane time constant and *I* (specified below) is a combination of external and recurrent inputs. To model spiking, we set a threshold θ (typically θ=0.1) such that when the membrane potential *x*_*i*_(*t*) > θ, *x*_*i*_(*t*) is reset to 0 and instantaneous spiking rate *r*_*i*_(*t*) is set to 1. Spiking-evoked input to postsynaptic neurons was modeled as a synaptic current with dynamics 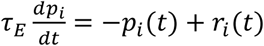, where *τ*_*E*_ is the decay time constant of excitatory currents. To simulate the slow excitatory currents produced by NMDA receptors, we set *τ*_*E*_ = 200 ms.

We next added recurrent connectivity between model units. Connectivity between model units is random and sparse, with p = 10% probability of a synapse forming between any two neurons, and weights of existing synapses sampled from a uniform distribution: 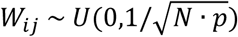. We also defined a gain parameter *g* that scales the strength of all synapses in the network.

To reduce finite-size effects in this model, we modeled recurrent inhibition by a single graded input *I*_*inh*_ representing an inhibitory population that receives equal input from, and provides equal input to, all excitatory units; dynamics of *I*_*inh*_ thus evolve as 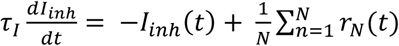, where *τ*_*I*_ = 50 ms is the decay time constant of inhibitory currents.

Each modeled “stimulus” input to the network was modeled with the same dynamics, with a high initial firing rate that decayed to a much lower sustained firing rate, and dropped to zero ten seconds after stimulus onset: specifically, in our model this input took the form 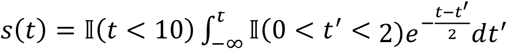 where 𝕀 is the indicator function. Each stimulus drove a random 50% of excitatory units in the network with input strength *w*_*i*_~ *g* ∙ *U*(0,1).

Thus, outside of spiking events, the membrane potential of neuron *i* evolves as 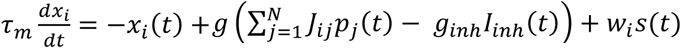. Model dynamics were simulated in discrete time using first-order Euler’s method with a timestep of dt = 1ms; a small Gaussian noise term *η*_*i*_~ 𝒩(0,1)/5 was added at each timestep. We explored model dynamics over a range of values of *g* and *g*_*inh*_, by selecting a value of *g* and performing a grid search over *g*_*inh*_ until the desired degree of persistence was achieved. Figures in the paper correspond to *g* = 1, *g*_*inh*_ = 3.8.

### Spiking recurrent neural network model + peptidergic excitation (pRNN)

In experimenting with the RNN+NMDA model described above, we found that we could achieve diverse temporal dynamics of spiking neurons if the time constant of excitation (*τ*_*E*_) was further increased, causing excitation to be much slower than inhibition. This allows model neurons to act as leaky integrators of their excitatory inputs, and start spiking when the population average activity (reflected by inhibitory input) drops below the integrated excitation. Because VMHdm^SF1^ neurons are known to express both glutamate and a variety of neuropeptides + neuropeptide receptors, we further modified the model by replacing the excitatory current with a mix of fast (glutamatergic) and slow (peptidergic) excitatory neurotransmission, although similar results are obtained in a model with just the slow component of excitatory neurotransmission.

We modeled fast excitatory currents as in the prior model, with dynamics 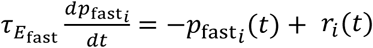, however we set 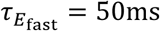 to better match the decay time constant of glutamatergic excitation. To model slow peptidergic excitation, we assumed that when a neuron spiked, peptide release was contingent on the recent firing rate history of that neuron, with peptide release only occurring if the average number of spikes in the last second exceeded a threshold *T* (typically *T* = 20, although performance was not strongly dependent on this parameter.) That is, the spiking of neuron *i* evoked peptide release if 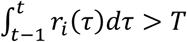. Dynamics of peptide-mediated excitation were otherwise modeled as before, thus giving 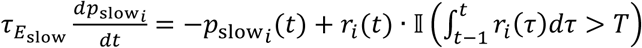, where 𝕀 is the indicator function. We used 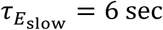 for all versions of the pRNN except for the third (black traces in Fig 4), for which 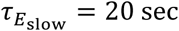 (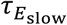 is abbreviated as *τ*_*S*_ in Fig 4).

For simplicity we assumed the synaptic weight matrix *J* was the same for both fast and slow components of excitation. Membrane potential dynamics in this model are therefore given by 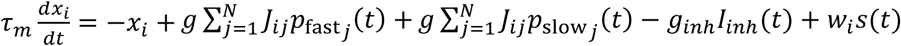. We present three versions of this model in Fig 4: in the “low gain” model, *g* = 1, *g*_*inh*_ = 8.8, 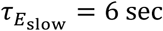; in the “high gain” model, *g* = 6, *g*_*inh*_ = 7.8, 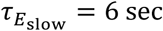; in the “high *τ*_*S*_ ” model, *g* = 2.5, *g*_*inh*_ = 4.25, 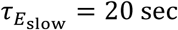. Simulation was performed as for the NMDA-RNN model, and as above parameters were fit by fixing the value of *g* (and 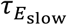) and performing a grid search over values of *g*_*inh*_ to achieve the desired degree of persistence.

### PRNN + local connectivity

The locally connected version of the pRNN model was created by adding a “distance dependence” on the probability of a pair of neurons forming a synaptic connection. Model neurons were numbered between 1 and N, and for neurons *i* and *j* the probability of forming a synapse was defined as 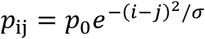, where *p*_0_ = 0.1 is the baseline degree of connectivity in the network, and σ sets the rate at which connectivity falls off with distance (here distance is defined as |*i*− *j*|). We found that broad connectivity was necessary to match the stimulus representation overlap seen in the data; plots in Fig 4 and the illustration of distance-dependent connectivity in **ED Fig 9a-b** were constructed using σ = 0.7*N*.

As in the pRNN, each stimulus in the local connectivity model provided input to 50% of model neurons. To match the observed Pearson’s correlation of the data, we found that it was necessary for stimulus inputs to reflect the structure of the model network, by targeting separate but still overlapping portions of the band of model neurons. Specifically, we found that the data was well fit when the middle 50% of model neurons in the band could receive input from both rat and USS stimuli, while the outermost 25% could only receive rat or USS input (see **ED Fig 9a**).

### Data similarity score, time-evolving dynamics

We constructed a data similarity score to quantify the degree of similarity between the plotted curves in Fig 4e, thus capturing how much the time-evolving dynamics of model neurons looked like that of the data. For each model and each mouse, we computed the Mean Correlation as defined above, which we will call *MC*_model_(*t*) for a given model and *MC*_mouse *i*_(*t*) for a given mouse. MC is a function of time--thus to quantify the mean similarity between the data and a given model over time, we considered the value of *MC*_model_(*t*) and *MC*_mouse *i*_(*t*) for all imaging frames (acquired at 11Hz) from 0 to 45 seconds relative to stimulus onset, which we reference using a frame index *t* = 1…T (so t=1 corresponds to a time of 0sec and t=T corresponds to a time of 45sec). Given these definitions, we define the data similarity score of the model dynamics as:

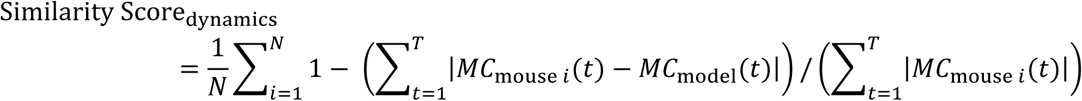

This can be simply interpreted as akin to the area between the data/model curves for each plot in Fig 4e. Note that the MC for the data here was computed from the USS-evoked neural activity, however MC for other stimuli gave similar results, as we found little difference between the MC for different stimuli.

### Data similarity score, stimulus specificity

This data similarity score quantifies the degree of similarity between the plotted curves in Fig 4h, ie how much the Pearson’s correlation between rat- and USS-evoked activity in each model looked like that observed in the data. We computed the Pearson’s correlation (as defined above) for each model and each mouse, which we call *PC*_model_(*t*) for a given model and *PC*_mouse *i*_(*t*) for a given mouse. We define frames *t*=1…T as all imaging frames from times 0 to 45 seconds relative to stimulus onset (same as for the similarity score of dynamics). We then define the data similarity score of model stimulus-specific activity as:

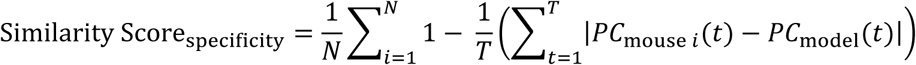

Like the similarity score of the dynamics, this can be interpreted as the area between the data/model curves for each plot in Fig 4h.

---

**Table 1:** Statistical significance testing. All t-tests are two-sided unless otherwise stated. All tests from distinct samples unless otherwise stated.

| Figure | Panel | Identifier | Sample size | p-value | Statistical test | Notes |
| --- | --- | --- | --- | --- | --- | --- |
| 1 | e | rat vs toy | 4 mice | 0.0078 | paired t-test |  |
|  | g | rat vs mouse | 8 mice | 0.0001 | paired t-test |  |
|  |  | rat vs toy | 8 mice | 0.0005 | paired t-test |  |
|  |  | mouse vs toy | 8 mice | 0.0015 | paired t-test |  |
|  | h | rat vs mouse - headfixed | 4 mice home cage, 8 mice head fixed | 0.0052 | paired t-test |  |
|  | i | rat vs mouse - headfixed | 4 mice home cage, 8 mice head fixed | 0.0110 | paired t-test |  |
|  | k | rat control vs rat iC++ | 7 mice control, 7 mice iC++ | 0.0289 | t-test |  |
|  | m | during+after vs control | 12 mice control, 6 mice during+after | 0.0078 | rep. meas. ANOVA | covariates tested: group vs time |
|  |  | after only vs control | 12 mice control, 6 mice after-only | 0.0321 | rep. meas. ANOVA | covariates tested: group vs time |
|  | n | control before vs after rat | 12 mice | 0.0006 | paired t-test |  |
|  |  | control before rat vs after PS off | 12 mice | 0.0074 | paired t-test |  |
|  |  | after rat, control vs during+after | 12 mice control, 6 mice during+after | 0.0023 | t-test |  |
|  |  | after PS off, control vs during+after | 12 mice control, 6 mice during+after | 0.0223 | t-test |  |
|  |  | after PS off, control vs after only | 12 mice control, 6 mice after-only | 0.0062 | t-test |  |
|  | o | control vs during+after | 12 mice control, 6 mice during+after | 0.0003 | rep. meas. ANOVA | covariates tested: group vs time |
|  |  | control vs after only | 12 mice control, 6 mice after-only | 0.0044 | rep. meas. ANOVA | covariates tested: group vs time |
| 2 | d | * | 5 mice | 0.0117 | paired t-test |  |

**Extended Data 1.**
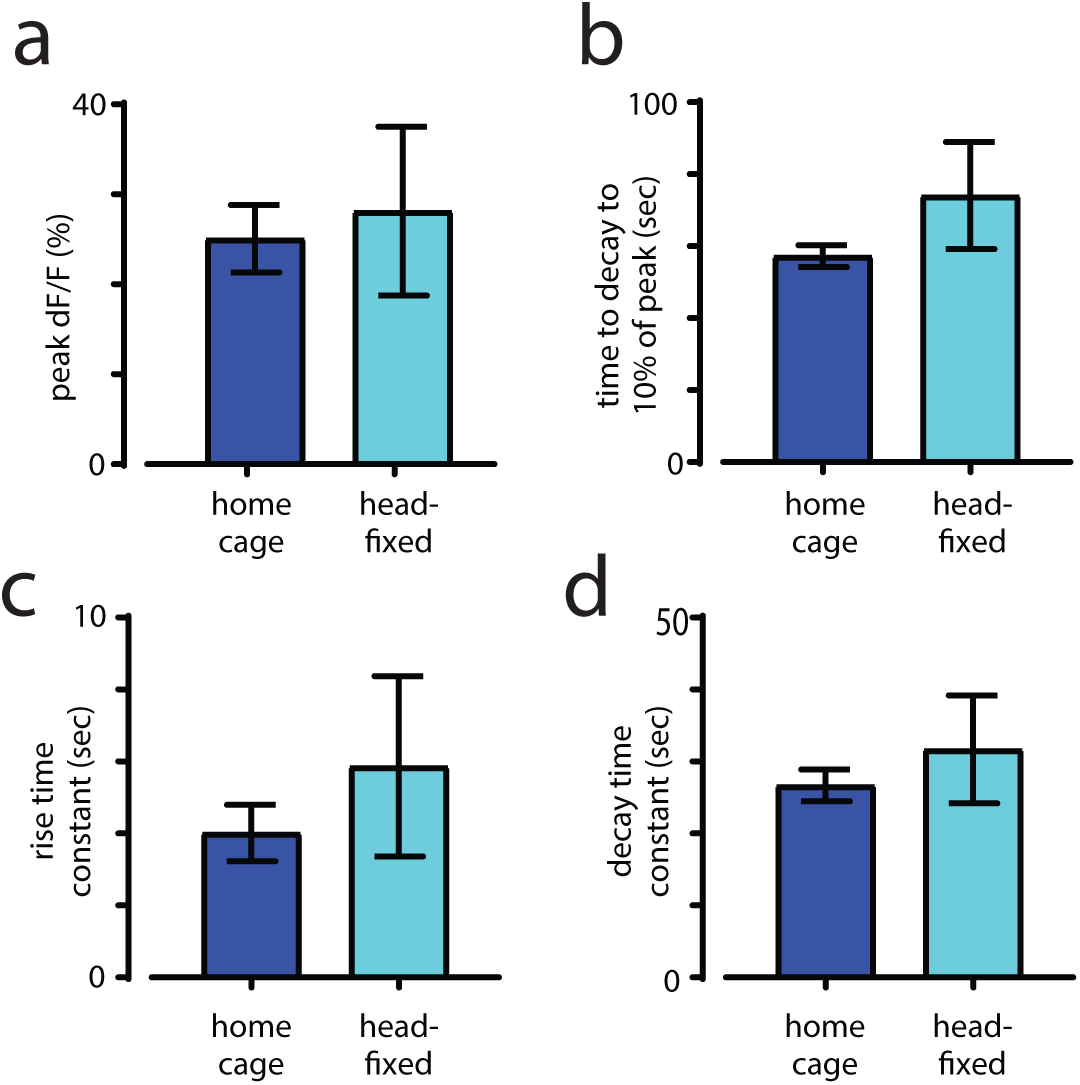
Comparison of SF1+ neurons’ response to rat in mouse home cage and head-fixed set-up. **a**, Peak ∆F/F activity in response to rat in home cage and a head-fixed set-up. (home cage group n = 4; head-fixed group n = 8). **b**, Response time measured at 10% of peak. **c**, Rise time constant measured as time elapsed to reach 1/e of peak amplitude. **d**, Decay time constant measured as time elapsed from peak to 1/e of peak.

**Extended Data 2.**
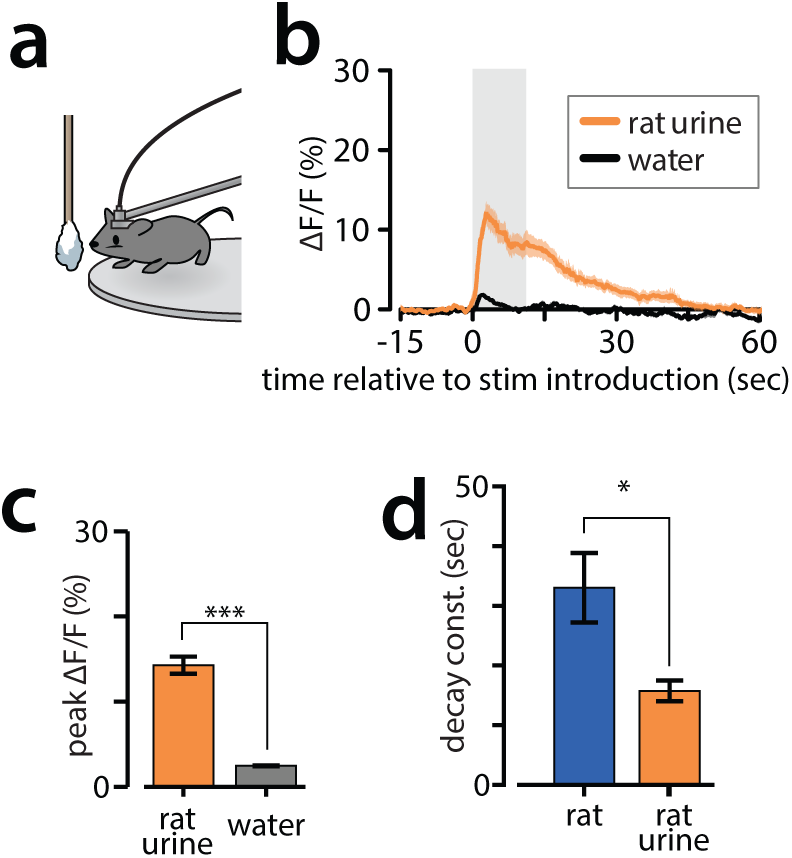
SF1+ neurons’ response to rat urine. **a**, Urine presentation to head-fixed fiber mouse. **b**, Averaged ∆F/F activity traces of SF1+ neurons in response to rat urine and water. n = 6. **c**, Peak ∆F/F activity triggered by rat urine and water. Paired t-test, n = 6. **d**, Decay time constant for rat and rat urine response, n = 6.

**Extended Data 3.**
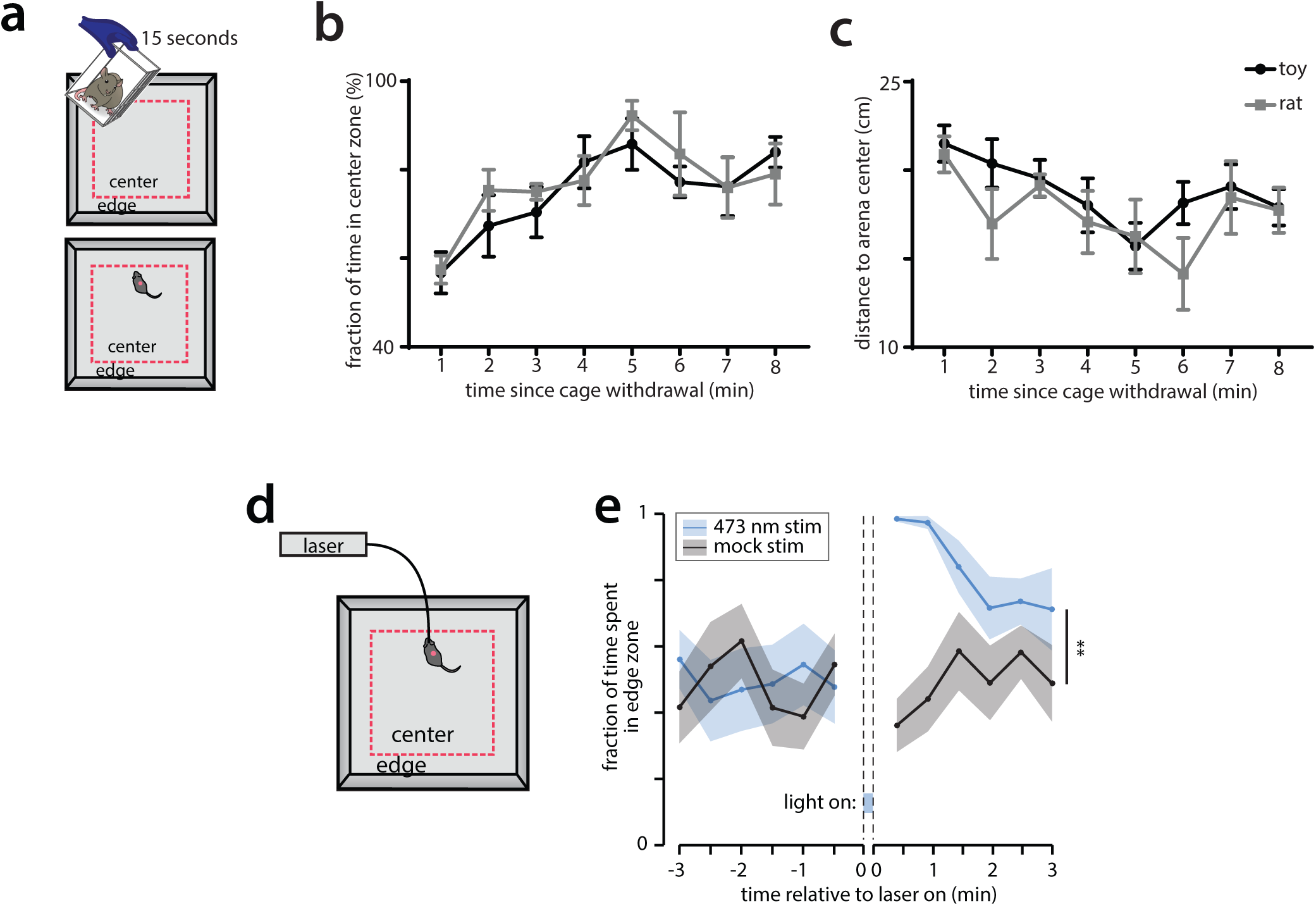
No change in mouse behavior due to potential lingering odor from rat. **a**, Schematic plot showing the experiment protocol: top, a live rat or toy rat (control) was brought to the open field arena in a wire mesh cage for 15 seconds; bottom, mouse was introduced to arena afterwards immediately. **b**, Fraction of time spent in center zone (as shown by red dashed line in a) for rat group and control group, n = 6 for each group. **c**, Distance from mouse body center to arena center. **d**, Schematic plot showing the optogenetic activation experiment protocol: mice expressing ChR2 was brought to the open field arena. After a 5 minutes habituation period, a 10 seconds light or mock stimulation was delivered to the mice. **e**, Fraction of time spent in the edge zone, n= 4 mice for each group.

**Extended Data 4.**
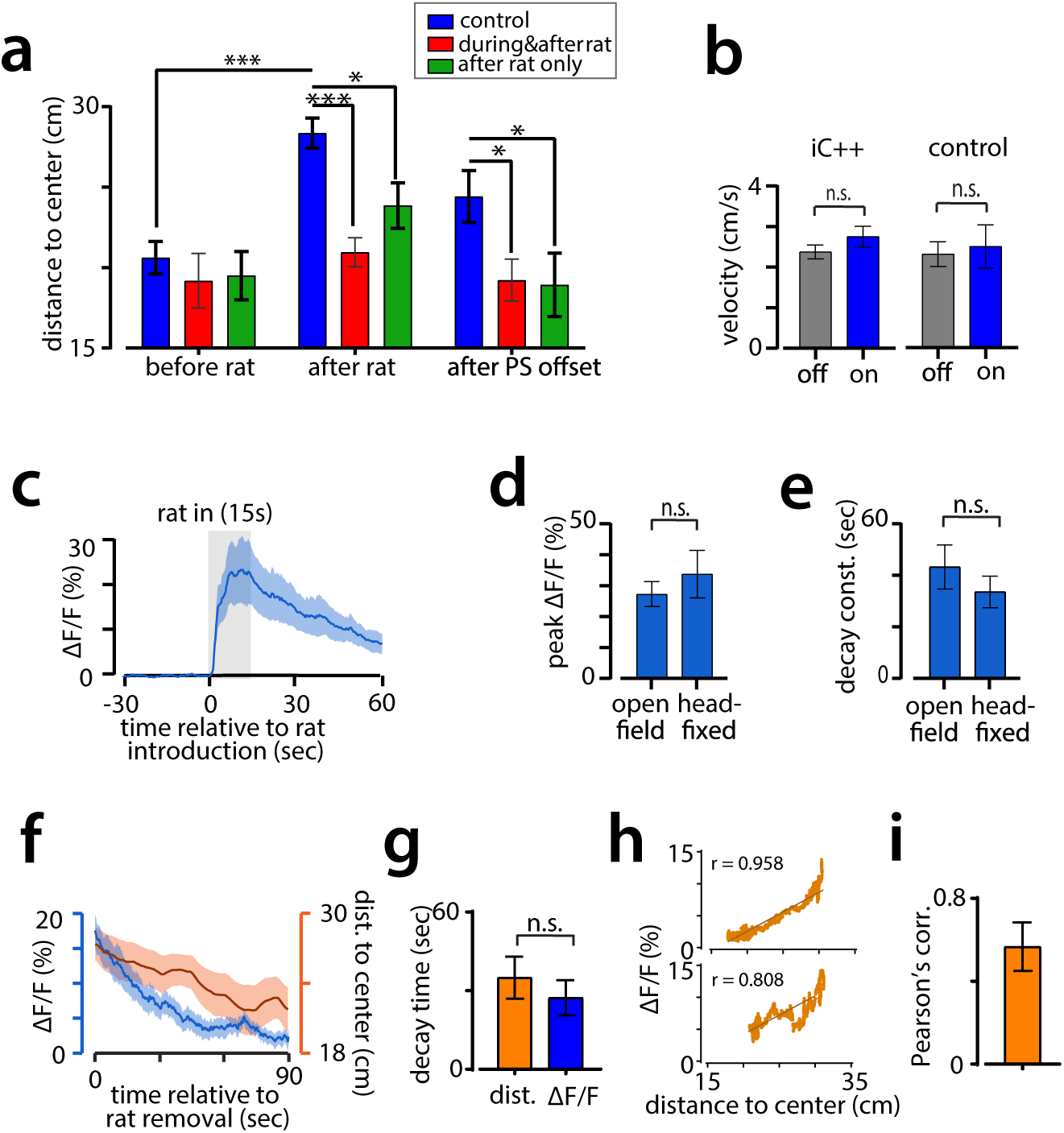
Fiber photometry and SF1+ neuron silencing in open field rat exposure assay. **a**, Distance from mouse body center to arena center during three different time periods: before rat, after rat and after photo stimulation offset, corresponding to −1~0, 0~1 and 3~4 minute in Fig.1m. **b**, Mean velocity was not altered by photo stimulation for iC++ and control mice. Velocity was measured in mouse home cage and averaged during a three-minute period for light off and light on sessions. Paired t test, n = 12. **c**, ∆F/F activity traces (mean ± SEM) of SF1+ neurons in response to rat in open field arena. Shaded gray bar denotes the presentation of rat. n = 9. **d**, Peak ∆F/F activity triggered by rat in open field arena (n = 9) and head-fixed set up (n = 8) (mean ± SEM). **e**, Decay constants of ∆F/F activity in open filed arena and head-fixed set up. **f**, Traces (mean ± SEM) for ∆F/F activity and the distance from mouse body center to arena center, aligned to rat removal. n = 9. Distance to center is plotted as 30 seconds moving average. **g**, Decay time measured as the time elapsed to reach 50% of the peak for linearly fitted data. n = 9, paired t test. **h**, Scatter plot of ∆F/F activity and the distance from mouse to arena center with a linear regression fitting for 2 example mice. Top, r = 0.958, p < 0.0001; bottom, r = 0.808, p < 0.0001. **i**, Pearson’s correlation coefficient between ∆F/F activity and the distance to center. n = 9.

**Extended Data 5.**
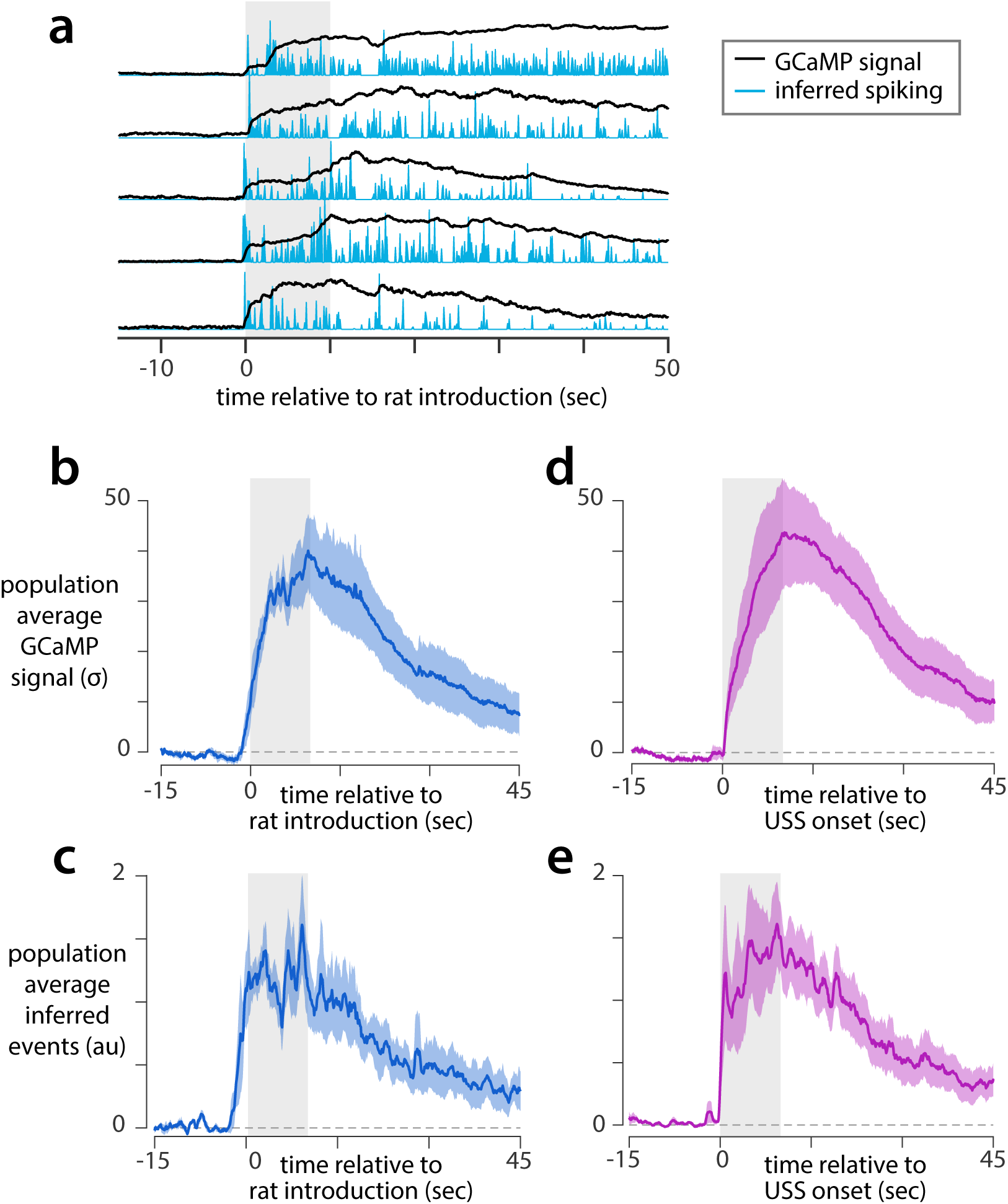
Inferred spiking of SF1+ neurons. **a**, Calcium traces (black) and spiking events inferred using constrained deconvolution1 (blue) for microendoscopic imaging of an example SF1+ neuron in response to five rat presentation trials in a head-fixed mouse. **b**, Response to one presentation of the rat stimulus in microendoscopic imaging experiments, averaged across cells (n=5 mice, mean ± SEM). **c**, Spiking events inferred from all individual SF1+ cells from the data in (b), averaged across cells and smoothed with a 1-second sliding window (n=5 mice, mean ± SEM). **d-e**, As in b-c, but for the USS stimulus.

**Extended Data 6.**
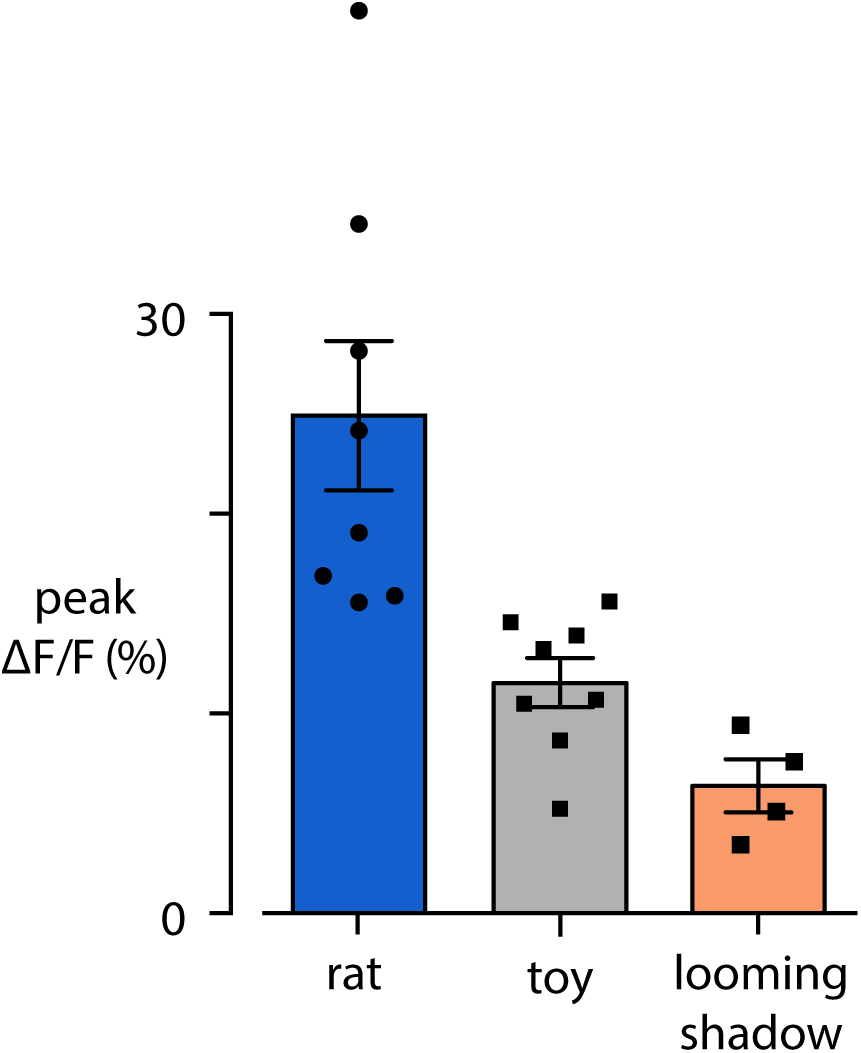
SF1+ population shows minimal response to the looming disk stimulus. Peak ∆F/F in response to rat, toy, or looming disk stimuli presented for 10 seconds in the animal’s home cage (points are individual mice).

**Extended Data 7.**
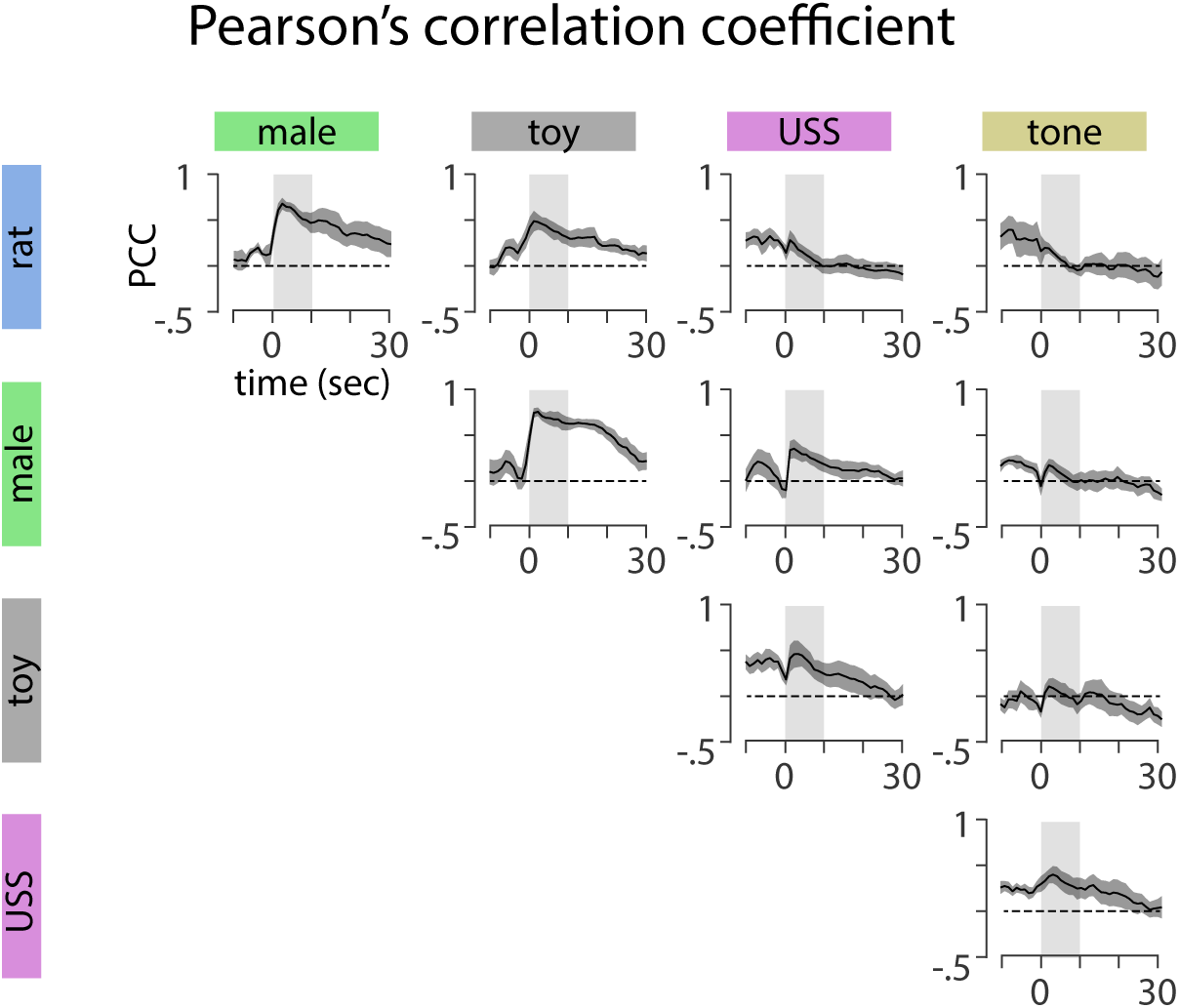
Additional Pearson’s correlations between stimulus pairs. Pearson’s correlation between SF1+ population activity evoked by all possible pairs of stimuli, as a function of time.

**Extended Data 8.**
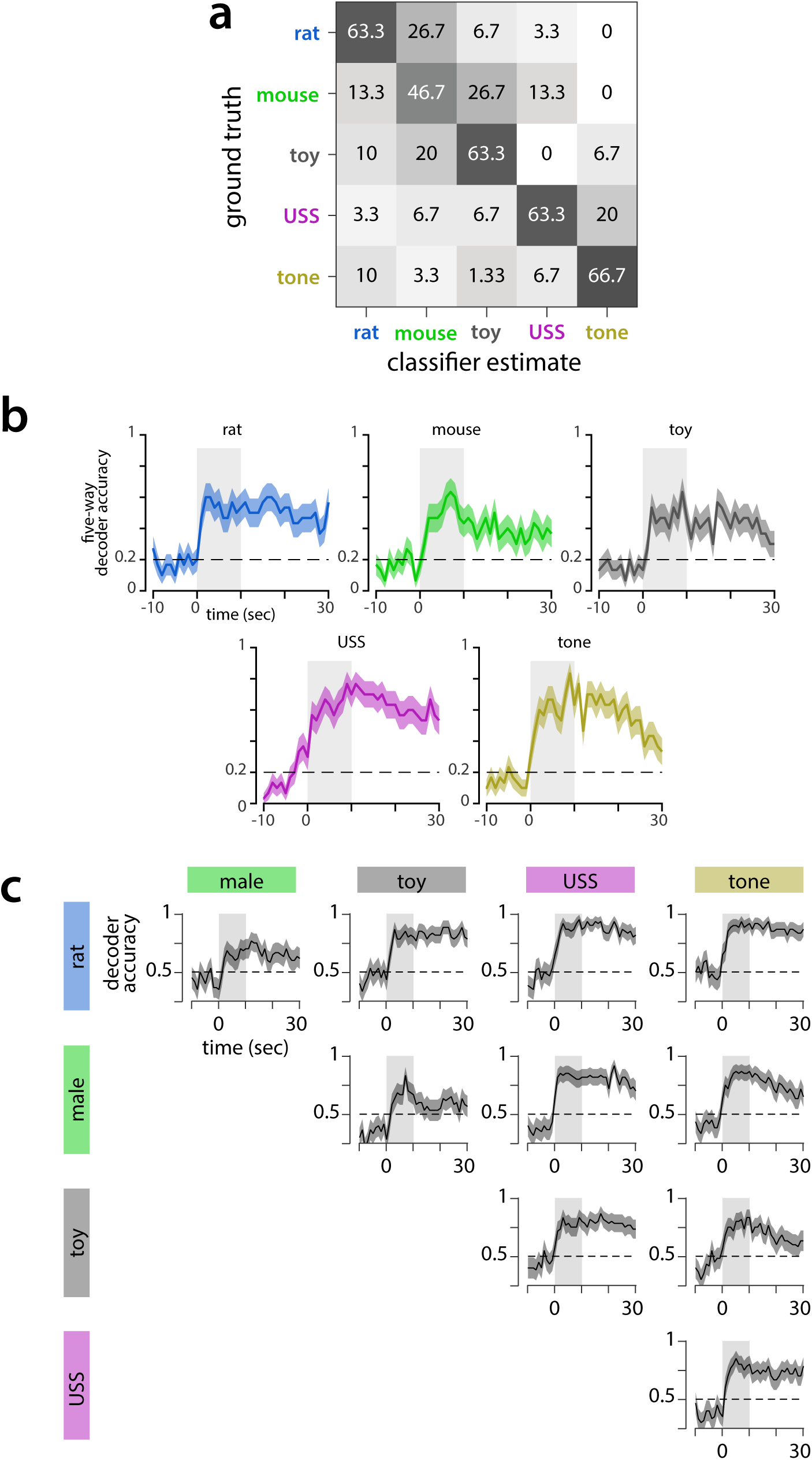
Additional decoder analysis of SF1+ population activity. **a**, Confusion matrix of the five-way Naïve Bayes decoder shown in Fig 3n, showing predicted stimulus identity for each stimulus class. Matrix is normalized so rows sum to 100%. **b**, Accuracy of the time-dependent five-way Naïve Bayes decoder shown in Fig 3o, as a function of time, for each tested stimulus. **c**, Accuracy of time-dependent binary Naïve Bayes decoders trained on all possible pairs of stimuli. The pair of stimuli being decoded for each plot is specified by the labels on the left and top. All plots show mean ± SEM across 5 imaged mice.

**Extended Data 9.**
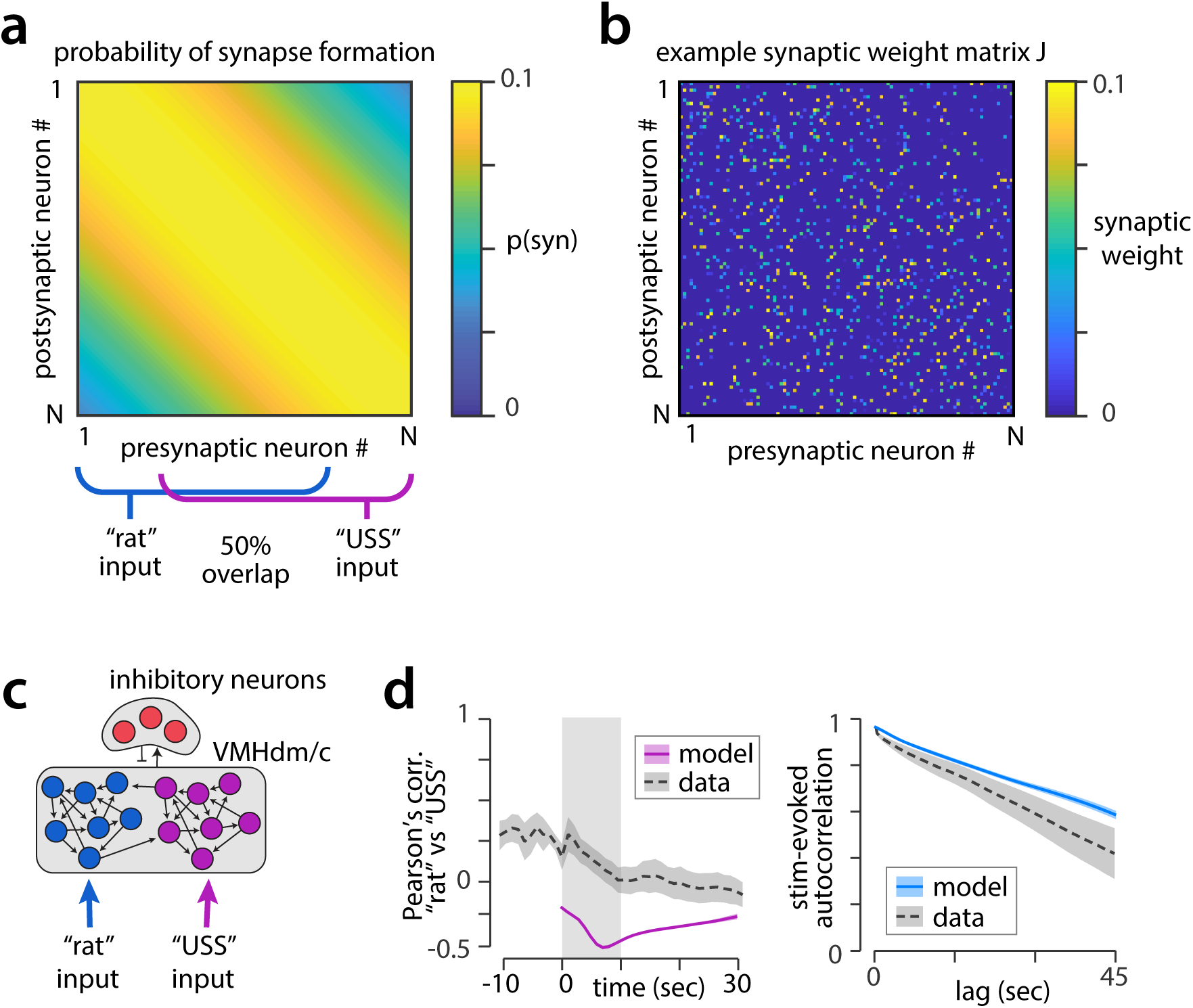
Locally connected model networks. **a**, Probability of synapse formation between neuron pairs decreases moderately as a function of “distance” (neuron number) in the locally connected sRNN model. Segments of the model targeted by rat and USS model input are also shown (blue/purple lines.) **b**, Example synaptic weight matrix generated from probability matrix shown in a; for visibility every 10th model neuron is shown. **c**, Example of a more highly structured model network, in which largely separate populations of neurons respond to the rat vs USS model inputs. **d**, Pearson’s correlation and stimulus-evoked autocorrelation for a network model such as that in (c), in which network structure results in no overlap between rat and USS representations.

